# Single-nucleus Transcriptomics Reveals Precystic Dysfunction in Polycystic Kidney Disease

**DOI:** 10.64898/2026.06.23.730438

**Authors:** Jonathan Marquez, Oleksandra Tymchyshyna, Sean K Gombart, Scott R Houghtaling, Grace Huang, Amrei M Mandel, Elizabeth D Nguyen, David R Beier

**Author notes:** To whom correspondence should be addressed: Jonathan Marquez, Center for Developmental Biology and Regenerative Medicine, Seattle Children’s Research Institute, 1900 Ninth Ave. Seattle, WA 98101. These authors contributed equally to this work (Elizabeth D Nguyen, David R Beier).

## Abstract

Polycystic kidney disease (PKD) is the most common cause of end stage renal disease with a known genetic etiology. This disease is characterized by the progressive development and expansion of kidney cysts. While recent studies have shed light on cell types and states contributing to PKD progression following cyst formation, the biological processes at work prior to cyst formation are relatively unexplored. To better understand mechanisms contributing to cystogenesis, we analyze pre-cystic kidneys from *Pkd1^R3277C/R3277C^*mice across multiple early timepoints, generating a transcriptomic atlas of nearly 1 million single nucleus transcriptomes. Activation of a small subset of genes in a precystic signaling pathway drives changes in both the distal convoluted tubule and proximal tubule cells. This pathway overlaps with a recently described “failed repair” transcriptomic signature despite the lack of clear changes in tissue morphology at these early stages of nascent cystogenesis. We identify Creb5 as a critical driver for cystogenesis. This single cell transcriptomic analysis of nascent cystogenesis reveals previously unrecognized cellular signaling at the earliest assessed points in precystic kidneys and provides a foundation for the development of high definition early diagnostic and therapeutic approaches prior to observable cysts in PKD.

## Introduction

Multiple genetic etiologies of polycystic kidney disease (PKD) have been identified since the term “polycystic kidney” was first used to describe a specific renal pathology in 1888 (*1*). The most common form of PKD, autosomal dominant polycystic kidney disease (ADPKD), is a disease with an estimated worldwide prevalence of 1 in 1000 individuals (*2*). Cysts arise in kidney tissue with disrupted function of the proteins encoded by mutated *PKD1* or *PKD2*(*3*, *4*). This appears to result in dysregulation of a wide variety of signaling pathways within cystic tissue that includes mTOR, WNT, and Hippo pathways (*5–7*). Enlarging cysts compress and impinge upon surrounding kidney parenchyma ultimately leading to chronic kidney disease (CKD) and eventual kidney failure (*8*). It is thus no surprise that decades of investigation have centered around understanding PKD through assaying cystic tissue. Yet, while these efforts have led to attempts to modulate pathways that seem to orchestrate cyst biology, most of these have had little success in affecting clinical outcomes. Therefore, alternative approaches to understanding PKD are paramount.

Recent advances in single-cell analysis of cellular and molecular complexity in models of PKD have bolstered our knowledge of changes occurring in cystic kidneys (*9–11*). These efforts have identified markers of transcriptional dysregulation within cysts and even detected a cell state within the proximal tubule that has been proposed as evidence of “failed-repair” within the proximal tubule (*9*, *11*). Yet, these studies have focused on tissue that has already become cystic. We posited that crucial insights into the pathogenesis of kidney cysts could be gained through assessing kidneys in a model of PKD prior to cyst formation.

One challenge to effectively assess pre-cystic kidneys in a model of PKD is the rapid progression of cyst formation in many of the available models of PKD. This makes testing kidney tissue at multiple timepoints before cyst formation difficult. To address this hurdle, we turned to a well characterized mouse model of PKD that develops cysts slowly due to a homozygous hypomorphic allele of *Pkd1*. Kidneys of *Pkd1^R3277C/R3277C^* (RC) mice remain largely devoid of cysts over the first month of life but consistently develop cysts over the course of the next few months (*12*). This contrasts with the rapidly developing cysts observed over the course of a few days or weeks observed in many of the widely used inducible models (*13*). This approach allowed us to assess for cell-type-specific expression signatures at pre-cystic timepoints in PKD.

We sought to work backwards in developmental time beginning at post-natal day 20 (P20), the earliest point where cysts can rarely be observed within kidneys of RC mice. We then assessed P10 and finally P5 to assess how early differing cell states could be detected in large single nucleus transcriptomic atlases of RC mouse kidneys. Through this approach we identified atypical cell signaling in the connecting tubule (CNT) and proximal tubule (PT) cells that correlate with the development of cyst-permissive tissue and are apparent as early as P10. Intriguingly these cells demonstrated an expression pattern similar to what has recently been described in multiple models of diverse kidney diseases as a “failed-repair” pathway (*9*, *14–17*), despite seemingly no morphological evidence of disrepair at the time points we assessed. We therefore propose that this expression pattern may define a fundamental process across kidney disease types and can occur early in disease without overt morphological damage to be repaired. We thus term the cells exhibiting this expression pattern: sentinel kidney disease (SKiD) cells. Our findings through analysis of single nucleus transcriptomic atlases will be the foundation to better understand the mechanism of pre-cystic PKD and develop therapeutic approaches to prevent cystic disease.

## Results

### Single-Nucleus Transcriptomic Profiling of Postnatal Pre-cystic PKD Kidneys

We sought to determine the earliest identifiable changes occurring in precystic PKD kidneys. To this end, we interrogated kidney-wide transcriptome changes in a stepwise fashion working backwards from the earliest morphological signs of disease. P20 kidneys display occasional small cystic changes, thus we chose this as the upper temporal bound of our inquiry. We performed snRNA-seq on four PKD and four control kidneys at P20 via sci-RNA-seq3 (*18*) (**Fig. 1A**). Each kidney was obtained from a distinct mouse. There were no discernable differences in size or weight between PKD and control kidneys at this stage (**Dataset 1**), though histological assessment of sections of the contralateral kidneys did show the expected limited morphological cystic changes (**Supplementary Fig. 1**).

**Figure 1:**
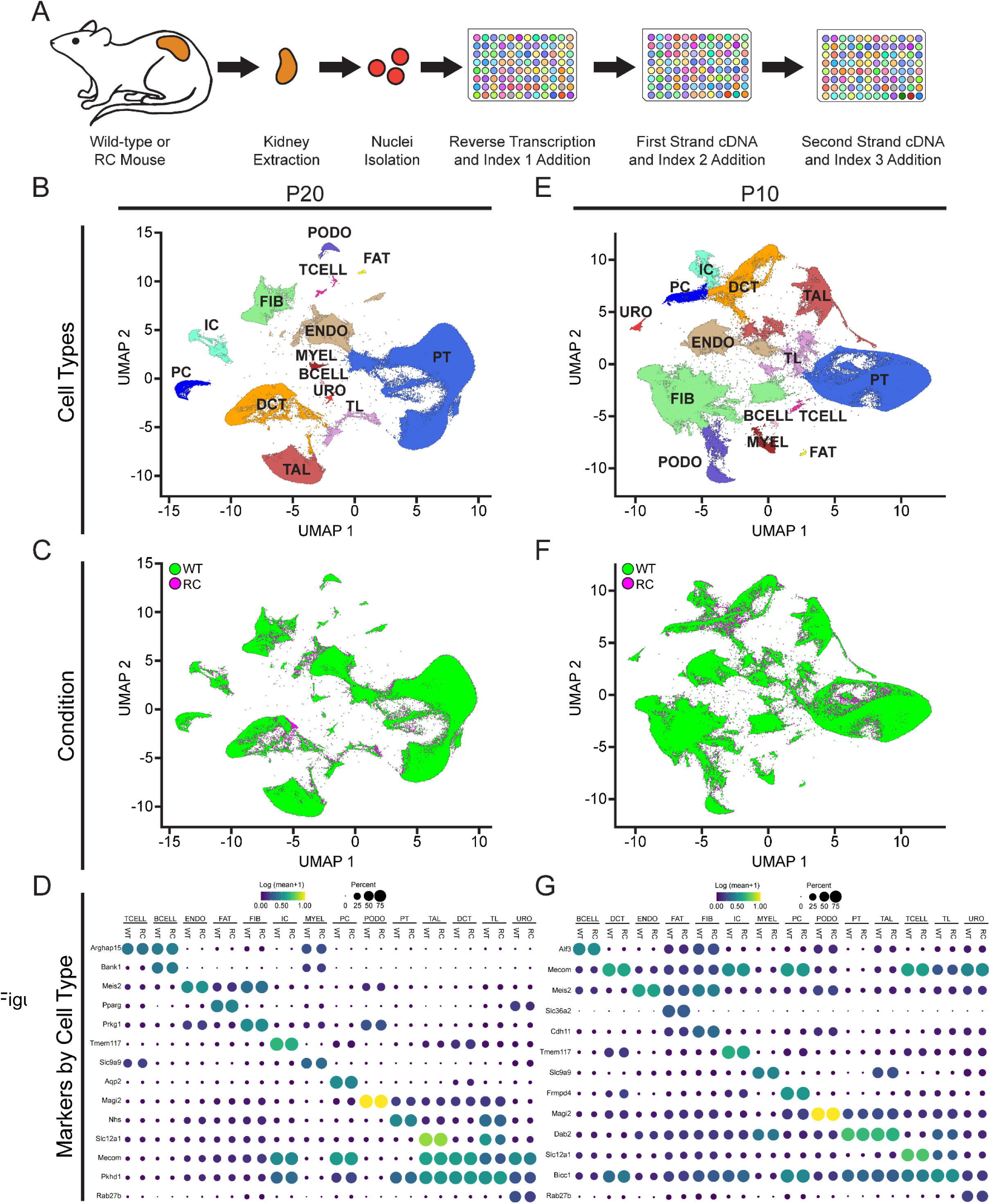
Transcriptomic Profiling of P20 and P10 Kidneys. (**A**) Overview of experimental approach. Combinatorial indexing was used to generate single-nucleus transcriptomic atlases from WT or PKD mouse kidneys. (**B**) UMAP plot of snRNA-seq dataset of combined P20 WT (n=4) and P20 RC (n=4) colored by cluster-based annotation. (**C**) UMAP plot of snRNA-seq dataset of combined P20 WT (n=4) and P20 RC (n=4) colored by tissue of origin genotype. (**D**) Dot plot of snRNA-seq dataset showing gene expression patterns of cluster-enriched markers for WT or PKD kidneys at P20. The dot diameter corresponds to the proportion of nuclei with detected activity of indicated gene, and the dot color corresponds to average gene activity relative to all nuclei. (**E**) UMAP plot of snRNA-seq dataset of combined P10 WT (n=3) and P10 RC (n=3) colored by cluster-based annotation. (**F**) UMAP plot of snRNA-seq dataset of combined P10 WT (n=3) and P10 RC (n=3) colored by tissue of origin genotype. (**G**) Dot plot of snRNA-seq dataset showing gene expression patterns of cluster-enriched markers for WT or PKD kidneys at P10. The dot diameter corresponds to the proportion of nuclei with detected activity of indicated gene, and the dot color corresponds to average gene activity relative to all nuclei. DCT: distal convoluted tubule cells; ENDO: endothelial cells; FIBRO: fibroblasts; IC: intercalated cells; PC: principle cells; PODO: podocytes; PT: proximal tubule cells; TAL: thick ascending limb cells; TL: thin limb cells; URO: urothelial cells.

After batch quality control filtering and preprocessing, PKD and control snRNA-seq datasets were integrated using Monocle(*19*) and visualized in UMAP space to annotate cell clusters. After removal of doublets and low-quality cells, we obtained a total of 363,945 nuclei by snRNA-seq; 180,861 nuclei from PKD and 183,084 nuclei from control kidneys. All expected cell types were identified in both PKD and control datasets (**Fig. 1B, C**). Lineage marker expression was well preserved in PKD kidneys as might be expected given the minimal morphological changes seen in PKD kidneys at this stage (**Fig 1D**). We cataloged differentially expressed genes in PKD for each cell type and detected differentially expressed genes across multiple cell types (**Dataset 2**).

Heartened by our ability to generate a high-quality snRNA-seq dataset at this early postnatal kidney stage, we next completed this analysis at successively earlier timepoints choosing times halfway between the previous timepoint and immediately following birth (P0). As previous studies have used kidney snRNA-seq datasets to detect disease specific cell subtypes through subclustering analyses (*9*, *11*, *16*), we elected to use this as a metric to determine an endpoint to our progressively earlier snRNA-seq experiments. For successive timepoints we conducted subclustering analyses for each cell type to detect PKD-enriched cell subtypes with a threshold of at least 66% of cells in a subtype derived from PKD kidney cells. Such a cell subtype was detected only in subclustered DCT cells of P20 kidneys.

We repeated this process with three PKD and three control kidneys P10 and obtained a total of 338,631 nuclei by snRNA-seq; 167,546 nuclei from PKD and 171,085 nuclei from control kidneys. All expected cell types were again identified in both PKD and control datasets (**Fig. 1E, F**) with well-preserved lineage marker expression seen in PKD kidneys (**Fig 1G**). We cataloged differentially expressed genes in PKD for each cell type and detected differentially expressed genes across multiple cell types (**Dataset 2**). PKD-enriched cell subtypes were detected in both subclustered DCT cells and subclustered PT cells of P10 kidneys.

Next, we completed this analysis for three PKD and three control kidneys P5 and obtained a total of 267,211 nuclei by snRNA-seq; 144,139 nuclei from PKD and 123,072 nuclei from control kidneys. While annotation of expected cell types was largely feasible across both PKD and control datasets (**Fig. 2A, B**), clusters representing fat cells and T-cells were not detected in our control dataset. Interestingly, several cell types annotated at P5 represent earlier developmental time points that appear to persist at this early postnatal stage (**Fig. 2A**). Indeed, persistent expression of early markers of multipotency such as *Lhx1*, *Wnt4*, and *Meox1* are apparent across many cell types (**Fig. 2C**). No PKD-enriched cell subtypes were detected across cell types annotated in P5 kidneys.

**Figure 2:**
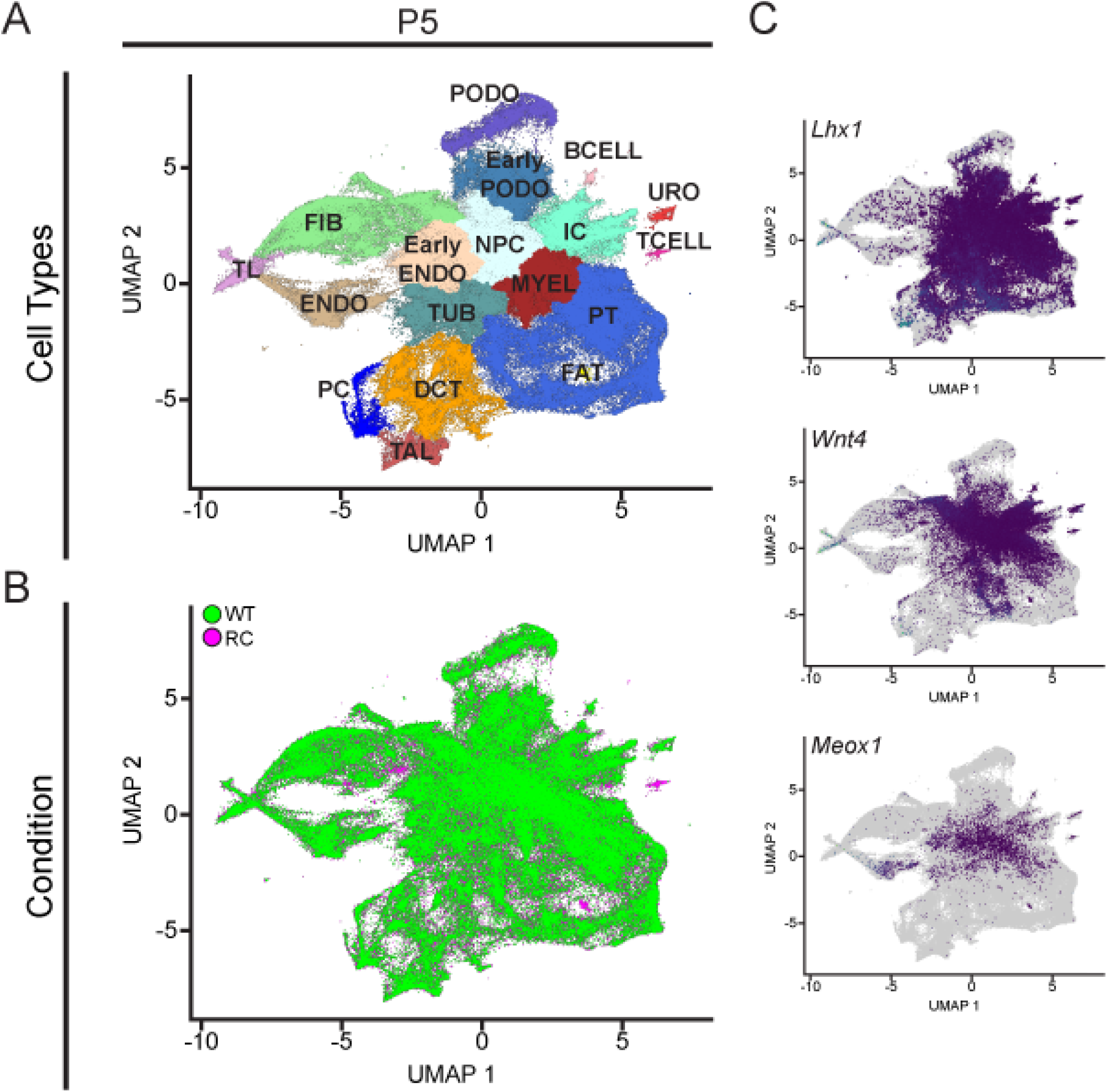
Transcriptomic Profiling of P5 Kidneys. (**A**) UMAP plot of snRNA-seq dataset of combined P5 WT (n=3) and P5 RC (n=3) colored by cluster-based annotation. (**B**) UMAP plot of snRNA-seq dataset of combined P5 WT (n=3) and P5 RC (n=3) colored by tissue of origin genotype. (**C**) UMAP plot of snRNA-seq dataset of combined P5 WT (n=3) and P5 RC (n=3) highlighting cells expressing early kidney cell markers *Lhx1*, *Wnt4*, and *Meox1*. DCT: distal convoluted tubule cells; ENDO: endothelial cells; FIBRO: fibroblasts; IC: intercalated cells; NPC: nephron progenitor cells; PC: principle cells; PODO: podocytes; PT: proximal tubule cells; TAL: thick ascending limb cells; TL: thin limb cells; TUB: unspecified early tubular cells; URO: urothelial cells.

### Molecular Pathways are Dysregulated in PKD Distal Convoluted Tubule Cells Across Early Postnatal Stages

To characterize PKD-enriched DCT cells, we annotated the subclustered DCT cells into cell subtypes based on marker genes and/or overrepresentation of cells derived from PKD kidneys. A threshold of 66% of the total number of cells derived from PKD kidneys was used to designate a cluster as PKD-enriched at each timepoint. At P20 subtypes of DCT and connecting tubule (CNT) cells were readily discernable (**Fig. 3A-C**). These subtypes were detected in approximately equal proportions in both PKD and control kidney cells (**Supplementary Fig. 2**). This subclustering also identified a cell subtype highly enriched in PKD derived cells (71%), suggesting a distinctive CNT cell state at P20 driven by PKD1 dysfunction.

**Figure 3:**
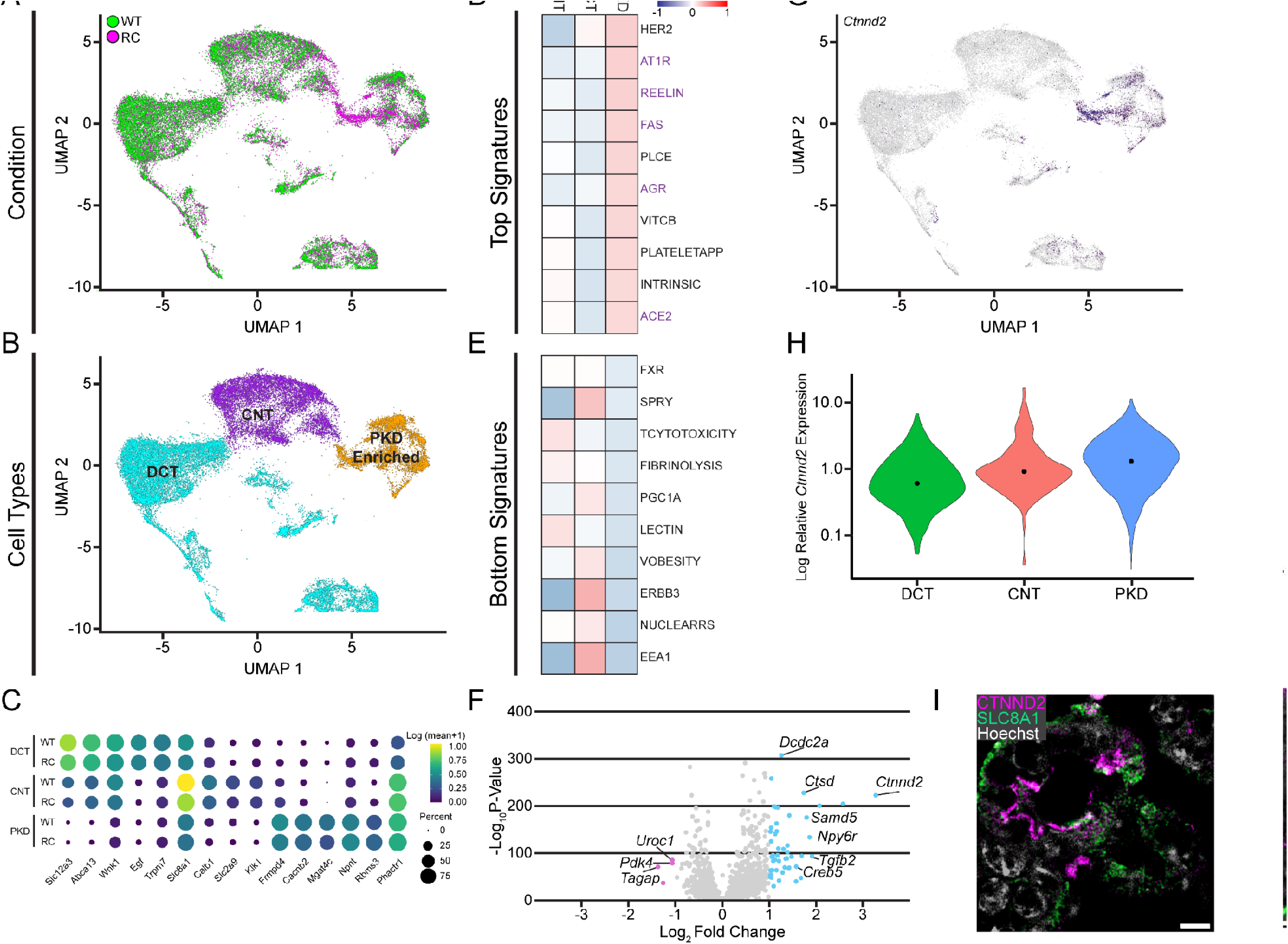
A PKD-enriched Cell State is Apparent in P20 CNT Cells. (**A**) UMAP plot of subclustered snRNA-seq dataset of P20 DCT cluster colored by tissue of origin genotype. (**B**) UMAP plot of subclustered snRNA-seq dataset of P20 DCT cluster colored by subcluster cell type annotation. (**C**) Dot plot of snRNA-seq P20 DCT cluster dataset showing gene expression patterns of subcluster-enriched markers for WT or PKD kidneys. The dot diameter corresponds to the proportion of nuclei with detected activity of indicated gene, and the dot color corresponds to average gene activity relative to all nuclei. (**D**) Heatmap showing the top 10 enriched BioCarta geneset signatures in the PKD-enriched subcluster compared to CNT and DCT subclusters. (**E**) Heatmap showing the bottom 10 enriched BioCarta geneset signatures in the PKD-enriched subcluster compared to CNT and DCT subclusters. (**F**) Volcano plot showing differentially expressed genes in the DCT cluster of PKD mice compared to WT at P20. The x-axis represents the log-fold change, and the y-axis represents the adjusted P-values. (**G**) UMAP plot of subclustered snRNA-seq dataset of P20 DCT cluster highlighting cells expressing *Ctnnd2*. (**H**) Violin plots of relative *Ctnnd2* gene expression between subclusters. (**I**) CTNND2 localizes to the apical aspect of CNT cells in PKD kidneys. Scale bar indicates 5 μm.

To better understand the molecular signature of this PKD-enriched CNT cell subtype, we applied pathway enrichment analysis across subtypes using VISION (*20*) (**Fig. 3D, E**). We observed activation of a cohort of pathways associated with cytoskeletal remodeling (**Fig. 3D, E pathways in purple text**) at P20, implicating these processes in a cell state that is permissive for cyst development. Certainly, cytoskeletal changes have been implicated in the pathogenesis of PKD and evidence of polycystin 1 as a mechanosensor has been suggested as a direct link to how its dysfunction may translate to these changes (*21–24*). In contrast, pathways related to energy metabolism and homeostasis appear diminished within this PKD-enriched subset, presumably reflecting dysregulated cellular metabolism that could precede cyst formation. This observation is in line with dysregulation of metabolism that has been described in later stages of PKD across human and mouse studies (*25–27*).

Differentially expressed genes observed within the DCT cluster at P20 (**Fig. 3F**) include genes that may participate in cytoskeletal remodeling as well as genes that may drive larger changes in PKD specific signaling. Among the differentially up-regulated genes in the DCT cluster, the most upregulated gene was *Ctnnd2* that encodes catenin delta-2, which is an adhesive junction associated protein of the armadillo/beta-catenin superfamily that promotes the disruption of E-cadherin-based adherens junctions to favor cell spreading upon stimulation by growth factors (*28–30*). Importantly, the nuclei expressing higher levels of *Ctnnd2* were within the subcluster of cells enriched in PKD (**Fig. 3G, H**). Fluorescence immune labeling analysis further identified increased CTNND2 in P20 PKD CNT cells (**Fig. 3I and Supplementary Fig. 3A**).

*Creb5* was the most differentially up-regulated gene encoding a transcription factor in PKD DCT cluster cells (**Fig. 3F**). The encoded cAMP responsive element-binding protein 5 cooperates with a variety of binding partners to coordinate gene expression (*31–34*). Intriguingly, this along with other upregulated PKD DCT cluster genes such as *Tgfb2* have been described as part of a pathway of failed repair in various kidney disease states (*9*, *11*, *14*, *16*, *35*). Yet, such a pathway had not been previously described as active without visible morphological or mechanical changes such as cysts or ischemia.

At P10 subtypes of DCT cells were also readily discernable (**Fig. 4A-C**), yet no CNT cell subtype could be distinguished independent of a PKD-enriched (71%) cluster. This potentially indicates that PKD-enriched cells at this timepoint are more alike to the typical CNT cells of the nephron at this stage. Yet, through pathway enrichment analysis we again saw the activation of pathways associated with cytoskeletal remodeling (**Fig. 4D pathways in purple text**) at P10. This suggests that signaling to modify cellular cytoskeleton is a longitudinal process occurring early in postnatal precystic kidney PKD pathogenesis. Similarly, pathways related to energy metabolism and homeostasis appear diminished within the PKD-enriched subset even at this earlier stage (**Fig. 4E**).

**Figure 4:**
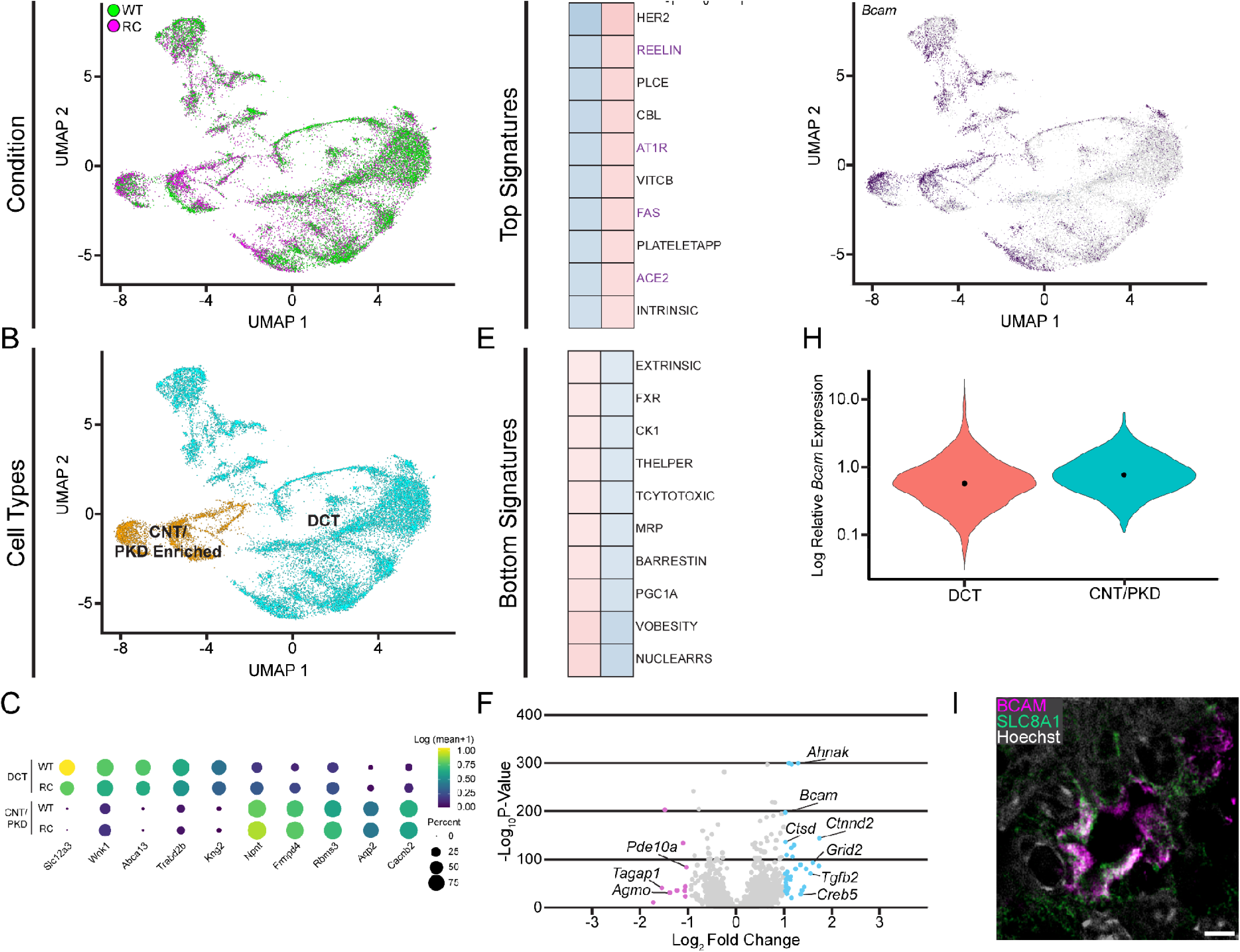
A PKD-enriched Cell State is Apparent in P10 CNT Cells. (**A**) UMAP plot of subclustered snRNA-seq dataset of P10 DCT cluster colored by tissue of origin genotype. (**B**) UMAP plot of subclustered snRNA-seq dataset of P10 DCT cluster colored by subcluster cell type annotation. (**C**) Dot plot of snRNA-seq P10 DCT cluster dataset showing gene expression patterns of subcluster-enriched markers for WT or PKD kidneys. The dot diameter corresponds to the proportion of nuclei with detected activity of indicated gene, and the dot color corresponds to average gene activity relative to all nuclei. (**D**) Heatmap showing the top 10 enriched BioCarta geneset signatures in the PKD-enriched subcluster compared to CNT and DCT subclusters. (**E**) Heatmap showing the bottom 10 enriched BioCarta geneset signatures in the PKD-enriched subcluster compared to the DCT subcluster. (**F**) Volcano plot showing differentially expressed genes in the DCT cluster of PKD mice compared to WT at P10. The x-axis represents the log-fold change, and the y-axis represents the adjusted P-values. (**G**) UMAP plot of subclustered snRNA-seq dataset of P10 DCT cluster highlighting cells expressing *Bcam*. (**H**) Violin plots of relative *Bcam* gene expression between subclusters. (**I**) BCAM localizes to the membrane of CNT cells in PKD kidneys. Scale bar indicates 5 μm.

There was a great deal of overlap between differentially expressed genes observed in the cells of the DCT cluster at P10 and P20. At P10 differentially expressed genes in DCT cluster cells (**Fig. 4F**) also included *Ctnnd2* and *Creb5*, further supporting the importance of cytoskeletal remodeling and the previously described repair associated pathway. Yet other genes such as *Bcam,* showed uniquely significantly increased expression only at this earlier timepoint. *Bcam* encodes the basal cell adhesion molecule that serves as a receptor and scaffold, which plays a crucial role in cell adhesion and motility (*36–38*). Importantly, the nuclei expressing higher levels of *Bcam* were within the subcluster of cells enriched in PKD (**Fig. 4G, H**). Fluorescence immune labeling analysis identified increased BCAM in P10 PKD CNT cells (**Fig. 4I and Supplementary Fig. 3B**). As with changes at P20, differential gene expression observed across the DCT cluster at P10 is driven by the increased expression of genes in the PKD-enriched CNT cell subtype (**Fig. 4F**). These findings collectively suggest that DCT cluster cell states show dynamic alterations in cytoskeletal remodeling amongst other signaling pathways during early precystic PKD pathogenesis.

As REELIN signaling was the most highly upregulated pathway in our analysis of both P20 and P10 PKD-enriched DCT subclusters, we sought to further evaluate the activity of this pathway. Yet, this was complicated by the observation that *Reln*, itself, was not transcriptionally upregulated at these timepoints (**Supplementary Fig. 4A**). As REELIN signaling has been described to occur independently of Reelin in a non-canonical or “inside-out” fashion (*39–42*), we examined the abundance of DAB1 which serves as the downstream effector of this pathway. Both DAB1 and the active phosphorylated form pDAB1 were increased in PKD kidneys (**Supplementary Fig. 4B**).

### Molecular Pathways are Dysregulated in PKD Proximal Tubule Cells in a Time Limited Fashion

To characterize PKD-enriched PT cells, we annotated the subclustered PT cells into cell subtypes based on marker genes and/or overrepresentation of cells derived from PKD kidneys with a threshold of 70% of cells at each timepoint. At P10 cell subtypes of proximal tubule segments 1-3 (PTS1, PTS2, and PTS3) were readily discernable (**Fig. 5A-C**). These subtypes were detected in approximately equal proportions in both PKD and control kidney cells (**Supplementary Fig. 5**). Subclustering also identified a cell subtype highly enriched in PKD derived cells (88%), suggesting a distinctive PT cell state at P10 driven by Pkd1 dysfunction. This appears to be unique to this timepoint of PT cells. Subclustering PT cells at P20, while readily classified into cell subtypes of PTS1, PTS, and PTS3 (**Supplementary Fig. 6**), did not reveal any subtype highly enriched in PKD derived cells.

**Figure 5:**
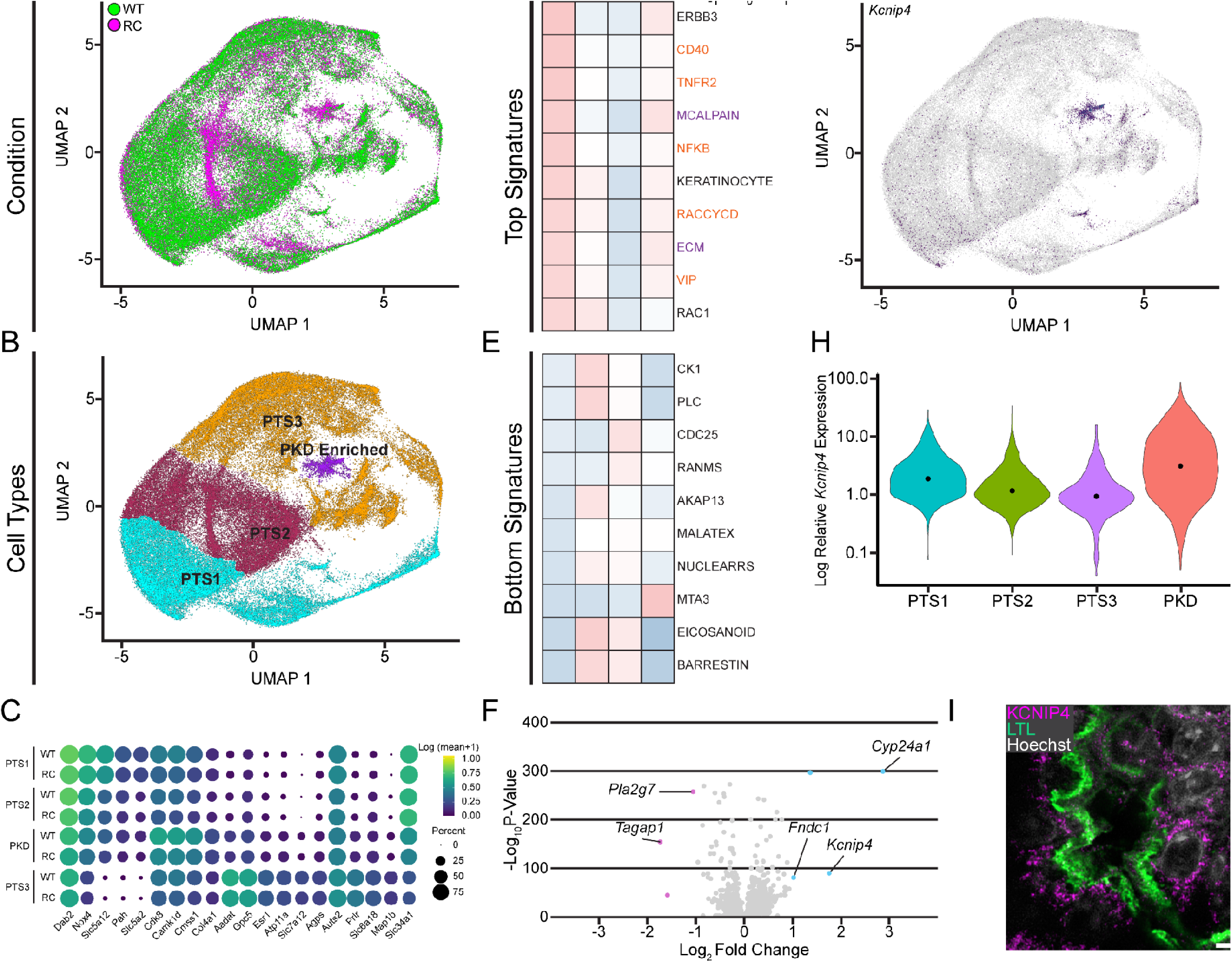
A PKD-enriched Cell State is Apparent in P10 PT Cells. (**A**) UMAP plot of subclustered snRNA-seq dataset of P10 PT cluster colored by tissue of origin genotype. (**B**) UMAP plot of subclustered snRNA-seq dataset of P10 PT cluster colored by subcluster cell type annotation. (**C**) Dot plot of snRNA-seq P10 PT cluster dataset showing gene expression patterns of subcluster-enriched markers for WT or PKD kidneys. The dot diameter corresponds to the proportion of nuclei with detected activity of indicated gene, and the dot color corresponds to average gene activity relative to all nuclei. (**D**) Heatmap showing the top 10 enriched BioCarta geneset signatures in the PKD-enriched subcluster compared to PTS1, PTS2, and PTS3 subclusters. (**E**) Heatmap showing the bottom 10 enriched BioCarta geneset signatures in the PKD-enriched subcluster compared to PTS1, PTS2, and PTS3 subclusters. (**F**) Volcano plot showing differentially expressed genes in the PT cluster of PKD mice compared to WT at P10. The x-axis represents the log-fold change, and the y-axis represents the adjusted P-values. (**G**) UMAP plot of subclustered snRNA-seq dataset of P10 PT cluster highlighting cells expressing *Kcnip4*. (**H**) Violin plots of relative *Kcnip4* gene expression between subclusters. (**I**) KCNIP4 localizes to PT cells in PKD kidneys. Scale bar indicates 5 μm.

Pathway enrichment analysis across PT subtypes at P10 also demonstrated activation of a smaller set of distinct pathways associated with cytoskeletal remodeling (**Fig. 5D, pathways in purple text**). Thus, distinct cytoskeletal remodeling may occur in the PT to allow for nephron morphology to move towards cyst formation. Yet, immune related pathways were also activated in this subset (**Fig. 5E, pathways in orange text**). This is consistent with the various studies that have implicated immune signaling in promoting or modifying the severity of cystic changes in PKD (*43–46*). Few pathways were downregulated in the PKD-enriched PT cell subtype (**Fig. 5E**) and for all these pathways, another cell subtype (i.e. PTS1, PTS2, or PTS3) had relatively lower levels of pathway activity.

The number of differentially expressed genes observed in PT cluster cells at P10 (**Fig. 5F**) are fewer in number than for the DCT cell cluster at P20 or P10. One of the most upregulated of these was *Kcnip4*, which encodes an inactivating cofactor that acts on a subset of potassium channels that facilitate cytoskeletal remodeling (*47*, *48*). Importantly, the nuclei expressing higher levels of *Kcnip4* were within the subcluster of cells enriched in PKD (**Fig. 5G, H**). Fluorescence immune labeling analysis identified increased KCNIP4 in P10 PKD PT cells (**Fig. 5I and Supplementary Fig. 3C**). These findings collectively suggest that PT cell states show dynamic alterations in cytoskeletal remodeling amongst other signaling pathways during early precystic PKD pathogenesis.

### Temporally Resolved Expression Patterns Facilitate Identification of Unifying Precystic PKD Drivers

With multiple timepoints of precystic PKD kidneys demonstrating signatures of possible disease processes in hand, we sought to integrate the temporal and cell subtype observations to determine the mechanisms driving precystic disease progression. While retaining cell subtype identities determined under initial analyses, we conducted unsupervised clustering of the combined cells of the P5, P10, and P20 DCT and PT cells. Supporting the robustness of the temporal context of cell states, cells clustered along both the lines of originally annotated cell type as well as with cells of their same timepoint (**Fig. 6A**). When the original cell subtype identity is projected onto the clustered cells, we observed that PKD-enriched cells were found in the cluster consisting of P10 PT as well as across P10 and P20 DCT (**Fig. 6A, B**, **dashed circles**). These temporal identities define a progression of transcriptional identities when assessed in trajectory analysis across pseudotime (**Fig. 6C**). This further establishes these cell subtypes as time limited in PT cells and persistent in DCT cells.

**Figure 6:**
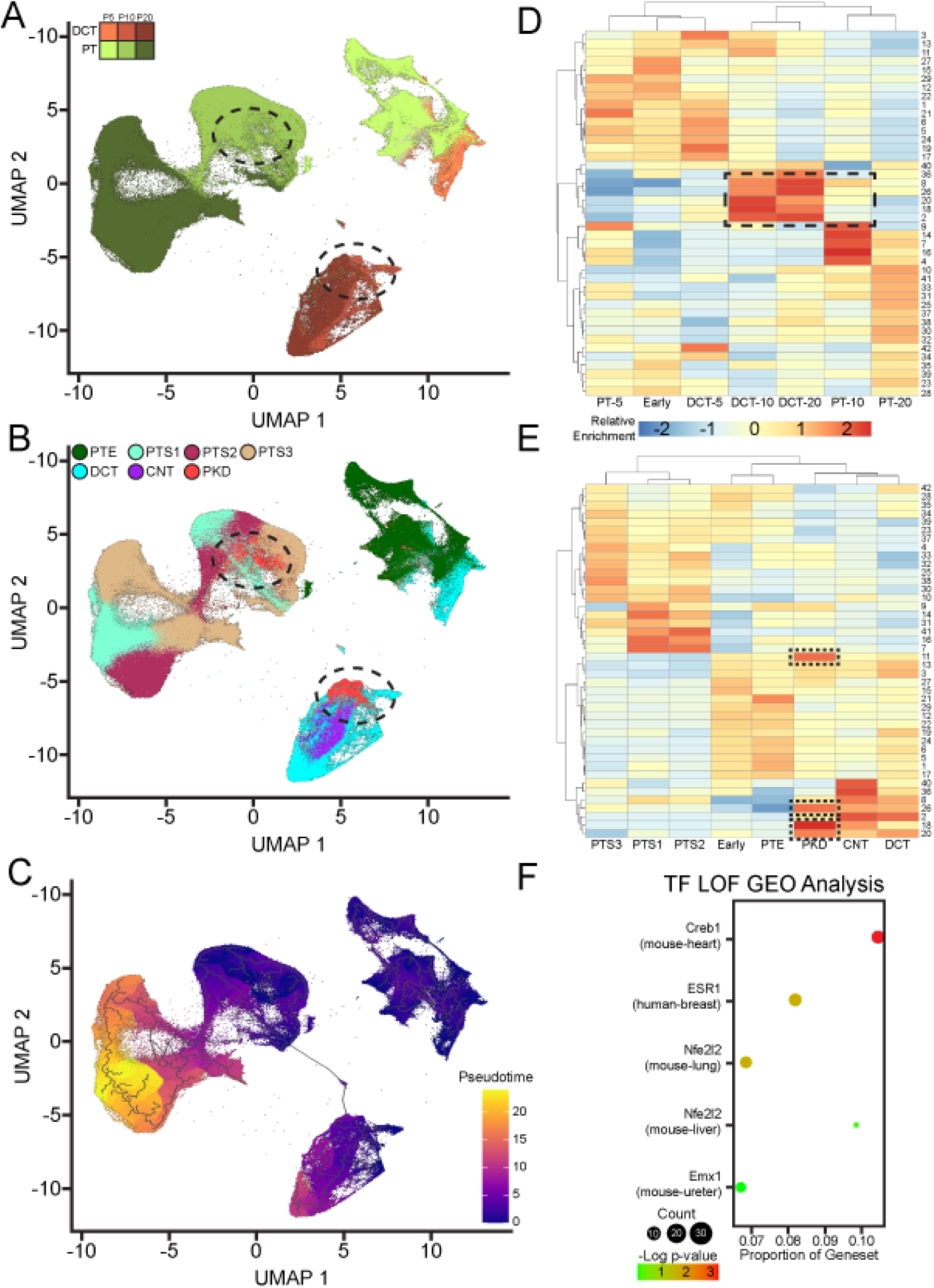
Temporal Analysis of Gene Expression in PKD-enriched Cell States Suggests a Role for CREB Signaling in Precystic PKD. (**A**) UMAP plot of combined PT and DCT clusters from P20, P10, and P5 WT and PKD kidneys colored by annotated cluster type in a temporal gradient. Dotted area contains the majority of cells from PKD-enriched subclusters. (**B**) UMAP plot of combined PT and DCT clusters from P20, P10, and P5 WT and PKD kidneys colored by annotated subclusters. Dotted area contains the majority of cells from PKD-enriched subclusters. (**C**) Pseudotemporal trajectory model superimposed upon the UMAP plot of combined PT and DCT clusters from P20, P10, and P5 WT and PKD kidneys colored by pseudotime. (**D**) Module analysis of coregulated gene expression patterns within the combined PT and DCT cluster nuclei from P20, P10, and P5 WT and PKD kidneys with modules distributed by cluster time. Dotted box indicates modules of genes that are most upregulated at time points that include PKD-enriched subclusters. (**E**) Module analysis of coregulated gene expression patterns within the combined PT and DCT cluster nuclei from P20, P10, and P5 WT and PKD kidneys with modules distributed by subcluster identity. Dotted box indicates modules of genes that are most upregulated within PKD-enriched subclusters. (**F**) Dot plot of enriched GEO genesets of response to transcription factor abrogation.

We hypothesized that we could use this information to understand precystic disease specific signaling independent of previously defined pathways. Further, in a parsimonious model, we hypothesized that a unifying signaling pattern could explain changes observed in both PKD-enriched DCT and PT cell subtypes. Such salient signaling would consist of groups of genes that would be increased in DCT cells at P10 and P20 compared to DCT cells of earlier timepoints and in PT cells at P10 compared to P5 and P20. We conducted gene module analysis which carries out UMAP space clustering with respect to gene expression, as opposed to with respect to cells as we had done previously. This identified 42 modules of seemingly coregulated genes (**Dataset 3**).

Mapping the modules onto cell type and timepoint identity identified 5 modules that were increased in DCT cells at P10 and P20 compared to earlier DCT cells and in PT cells at P10 compared to earlier or later PT cells (**Fig. 6C, dashed box**). We next mapped the same modules onto the cell subtypes that had been previously annotated to determine gene modules that were increased in PKD-enriched cell subtypes. This revealed 4 such modules (**Fig. 6D, dashed box**). As we posited that the most pivotal gene modules for PKD would be found by both these analyses, we next determined the overlap of modules identified by these assays and found 3 modules identified by both complementary approaches (**Supplementary Fig. 7**).

To analyze these high confidence gene modules, we next conducted gene set enrichment analysis to detect transcription factors that could be controlling expression of these gene modules. As we hoped to avoid the complications introduced by potential binding partners and varying chromatin states, rather than use CHIP-seq datasets to determine possible transcription factor activity, we turned to a dataset of experimentally derived single transcription factor loss-of-function and gain-of-function from across the Gene Expression Omnibus repository (*49*) that has been used to benchmark computational predictions of transcription factor activity (*50*). This analysis showed that the identified modules were enriched for genes with expression that decreased in response to murine *Creb1* loss-of-function (*51*) (**Fig. 6F**). Although we did not detect any evidence of increased *Creb1* expression in our precystic PKD analysis, *Creb5* expression was significantly elevated in PKD-enriched cells of the DCT cluster. Of note, CREB5 has been shown to share a binding site motif with CREB1(*52*, *53*).

### Subcellular Localization of PKD-enriched Transcription Factors Supports their Role in Driving Precystic Dysfunction

Given the potential importance of CREB signaling in PKD suggested by our transcriptome analysis, we sought to assess whether this was borne out at the protein level. No obvious differences were observed in CREB5 abundance between PKD and control kidneys at P5 through immunofluorescence labeling of endogenous Creb5 (**Fig. 7A**). Yet there was an increased abundance of CREB5 across CNT cells of PKD kidneys compared to controls in P10 and P20 kidneys (**Fig. 7B, C**). While this difference in Creb5 abundance was consistent with the increased transcription of *Creb5* observed in CNT cells within the transcriptomic atlas of PKD mice, we were struck by the dramatic increase in nuclear CREB5 observed in PKD CNT cells compared to control counterparts at P20 and P10 (**Fig. 7B, C**). This suggests that CREB5 is influencing precystic PKD changes through a nuclear role, likely in keeping with its known transcription factor function rather than some novel function unique to the kidney.

**Figure 7:**
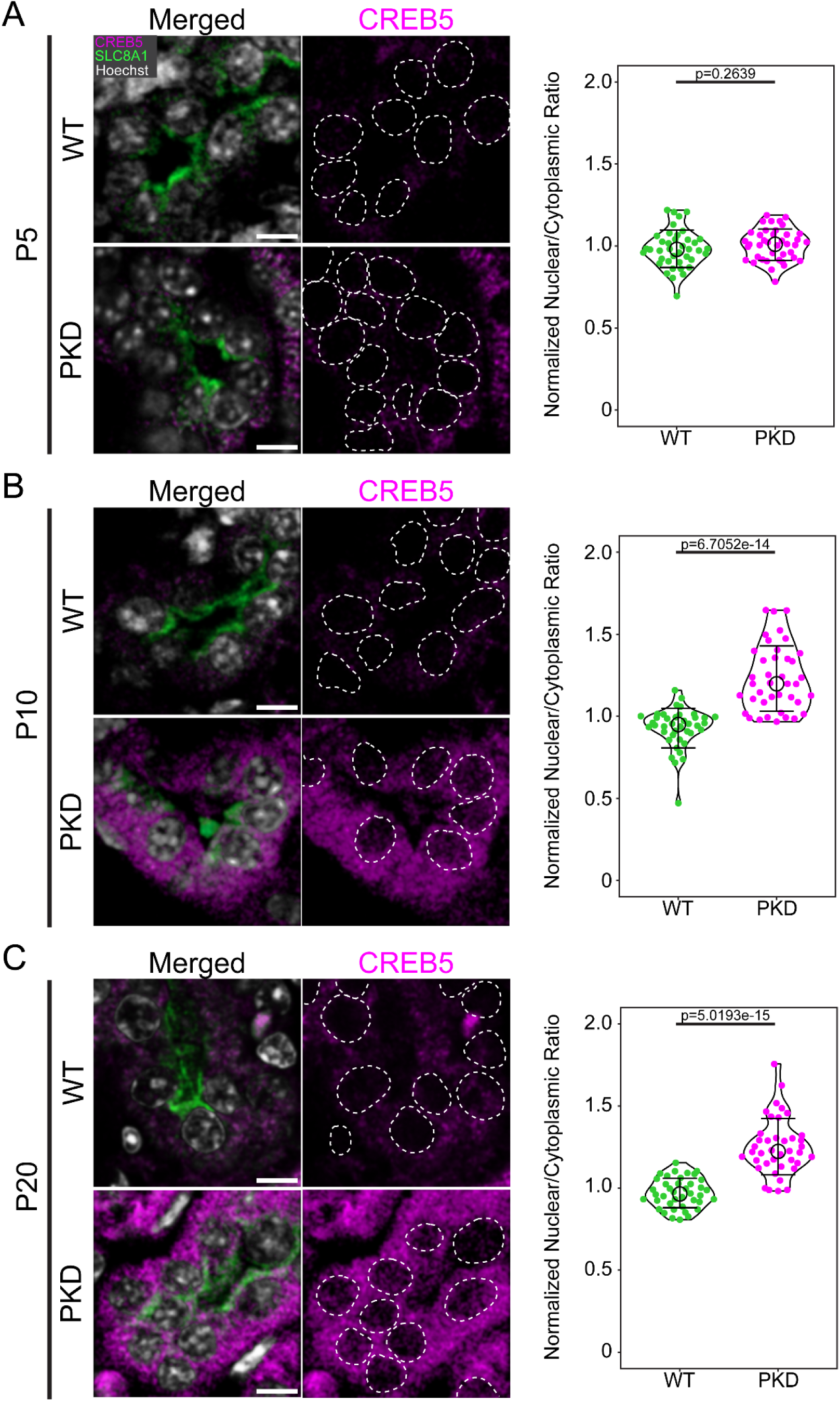
CREB5 Nuclear Localization is Increased in Precystic PKD CNT Cells. (**A**) Immunofluorescent labeling of CREB5 in WT (n=40) and PKD (n=40) CNT cells at P5 along with quantitation of normalized ratio of nuclear to cytoplasmic abundance. Dotted area indicates nuclei based on Hoechst staining. (**B**) Immunofluorescent labeling of CREB5 in WT (n=40) and PKD (n=40) CNT cells at P10 along with quantitation of normalized ratio of nuclear to cytoplasmic abundance. Dotted area indicates nuclei based on Hoechst staining. (**C**) Immunofluorescent labeling of CREB5 in WT (n=40) and PKD (n=40) CNT cells at P20 along with quantitation of normalized ratio of nuclear to cytoplasmic abundance. Dotted area indicates nuclei based on Hoechst staining. Dots are individual cell measurements, circles indicate the mean values, and whiskers indicate standard deviation. Statistical analysis was completed using two-tailed Mann Whitney Tests. Scale bars indicates 5 μm.

### Analysis of the Cell-Cell Communication Landscape Reveals Potential Biomarkers of Precystic PKD Progression

We sought to use analysis of cell-cell communication to identify signaling proteins that might be detectable as biomarkers of disease prior to cyst formation. To accomplish this, we used CellChat (*54*). We felt that the underlying database of ligands, receptors, and related cofactors used in this analysis of interactions would facilitate prioritization of signaling proteins that may be detectable extracellularly in clinically tractable specimens such as blood or urine rather than sequestered within kidney tissue.

Analysis of this cell-cell communication landscape readily identified differential strengths of interactions when comparing PKD to control kidneys across cell types at P10 (**Supplementary Fig. 8**). To focus our analysis on signaling that could be affecting the emergence of enriched cell states within the DCT and PT clusters at this stage we assessed differential strength of cell-cell communications within specific clusters to detect active communication which may be disease process-specific. This revealed that, for both DCT and PT clusters, the greatest differential cell-cell communication was seen in communication deriving from the PKD-enriched cell subclusters suggesting that the cells represented by the nuclei of these clusters are influencing their surrounding counterparts rather than being passive receivers of surrounding ligand-receptor mediated signaling (**Fig. 8A, B**).

**Figure 8:**
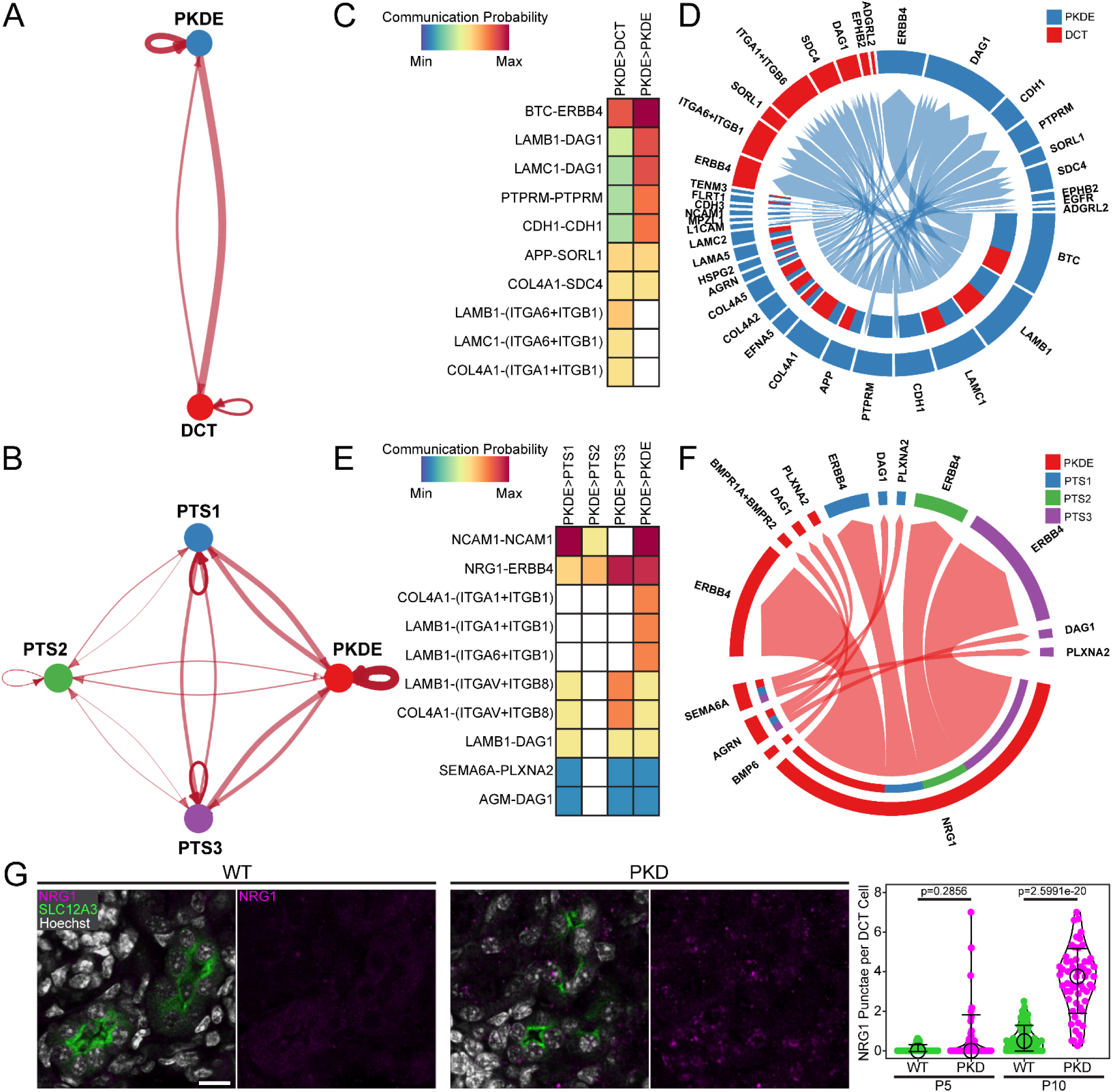
Precystic PKD Cell-Cell Communication Patterns Suggest Increased NRG Signaling that May Be Detectable in Urine. (**A**) String diagram of differential interaction strength between DCT and PKD-enriched subclusters between WT and PKD nuclei in the DCT cluster of P10 kidneys. Arrow width is proportional to increased communication in PKD tissue. (**B**) String diagram of differential interaction strength between PTS1, PTS2, PTS3, and PKD-enriched subclusters between WT and PKD nuclei in the DCT cluster of P10 kidneys. Arrow width is proportional to increased communication in PKD tissue. (**C**) Heatmap of changes in P10 DCT cluster cell-cell communication from sender PKD-enriched subcluster nuclei to all other subcluster nuclei in PKD tissue by ligand-receptor pair in PKD compared to WT nuclei. White boxes have no significant change in cell-cell communication probability. (**D**) Chord diagram of significantly increased cell-cell communication within the P10 DCT cluster in PKD compared to WT nuclei. Arrows connect ligand-receptor pairs. Colors indicate sender and receiver cell subcluster identities. (**E**) Heatmap of changes in P10 PT cluster cell-cell communication from sender PKD-enriched subcluster nuclei to all other subcluster nuclei in PKD tissue by ligand-receptor pair in PKD compared to WT nuclei. White boxes have no significant change in cell-cell communication probability. (**F**) Chord diagram of significantly increased cell-cell communication within the P10 PT cluster in PKD compared to WT nuclei. Arrows connect ligand-receptor pairs. Colors indicate sender and receiver cell subcluster identities. (**G**) Immunofluorescent labeling of NRG1 in WT (n=30) and PKD (n=30) tissue at P10 along with quantitation of abundance per DCT cell at both P5 and P10. Scale bars indicates 10 μm.

We assessed the composition of differentially increased ligand receptor pairs in both DCT and PT clusters. While a variety of ligand pairs with increased communication probability were evident, we were struck by the overlap of increased communication within NRG-ERBB signaling. Specifically, this was manifested as increased BTC-ERBB4 connectivity in signaling from PKD-enriched cells within the DCT cluster and as increased NRG1-ERBB4 connectivity in signaling from PKD-enriched cells of the PT cluster (**Fig. 8C-F**). Because this increased connectivity was observed within these clusters, we also assessed whether overall NRG signaling was enriched in our transcriptomic atlas at P10. The connectivity within the NRG pathway exists within the wild-type kidney and predominantly emanates from fat cells signaling to a variety of cell types (**Supplementary Fig. 9A**), commensurate with the role of NRG in adipose cell signaling (*55*, *56*). However, in PKD kidneys NRG signaling connectivity is significantly elevated from cells of thin limb of loop-of-Henle (TL) (**Supplementary Fig. 9B**). This increased TL NRG signaling is driven by signaling via the NRG1-ERBB4 ligand pair (**Supplementary Fig. 9C**) in PKD kidneys, compared to the predominance of signaling via the NRG4-ERBB4 ligand pair used by fat cells in both the PKD and wild-type kidneys (**Supplementary Fig. 9D**).

Given the properties of NRG1 as a cell surface protein that can be cleaved and serve as an extracellular signaling molecule (*57–59*), we were keen to evaluate this as a potential biomarker for precystic PKD disease progression. We assayed the kidneys of wild-type and PKD mice via immunofluorescence labeling and, while we saw no appreciable increase in NRG1 at P5 (**Fig. 8G**), we did observe a marked increase in NRG1 in P10 kidneys (**Fig. 8H**). While this was encouraging for the utility of NRG1 as a potential marker of precystic disease progression, we thought that it would be particularly useful as a biomarker if detectable in the urine as this is a particularly tractable sample type for laboratory testing. Evaluation of protein levels in the urine of wild-type and PKD mice showed detectable NRG1 in wild-type mice, and increased amounts in PKD mice (**Supplementary Fig. 10**).

## Discussion

Our study redefines our understanding of the temporal mechanistic origins of PKD, as early molecular events in *Pkd1* deficient kidneys arise well before the appearance of cysts. Through generating a single-nucleus transcriptomic atlas spanning nearly one million nuclei across three early postnatal stages, we show that cystogenesis is preceded by transcriptional changes in both distal and proximal nephron segments, despite preserved morphology. These findings support a model in which PKD is initiated not by a structural disturbance, but by a cell-intrinsic signaling program that prime the nephron for cystogenesis.

Our central discovery is that PKD-enriched cell states exist in CNT and PT cells prior to cyst formation. These cells exhibit a transcriptional program that overlaps with the “failed repair” state, yet they arise in the absence of disrupted morphology. This observation suggests that the signature of a “failed-repair” state is not exclusively a response to injury but may represent a sentinel program intrinsic to disease initiation. We therefore propose to designate these cells as sentinel kidney disease (SKiD) cells to reflect their early appearance and potential role as initiators of disease permissive tissue states.

Across nephron segments, SKiD cells appear to activate cytoskeletal remodeling pathways while downregulating a subset of metabolic programs. While cytoskeletal dysregulation has been implicated in cyst expansion, our data place these alterations at the earliest detectable stages of disease. Upregulation of genes such as *Ctnnd2* and *Bcam* suggest that factors such as junctional instability and altered mechanosensation could constitute foundational events that predispose tubules to dilate and ultimately form cysts. The transient appearance of PT SKiD cells at P10, contrasted with the persistence of CNT SKiD cells through P20, further suggests segment specific windows of vulnerability and potential for recovery that may shape cyst distribution and progression.

Integrative analysis of gene modules across timepoints suggested that changes common to SKiD cells across both PT and CNT cells may stem from CREB signaling as a unifying driver of early PKD pathogenesis. *Creb5* emerged as the most consistently upregulated CREB-family transcription factor across SKiD cells and its marked nuclear accumulation at P10 and P20 suggests an active regulatory role. Although previous work has identified increased expression of CREB target genes in PKD (*60*) and even increased expression of *CREB5/Creb5* in “failed repair” cells (*9*, *14*, *61*), our findings position CREB5 as a candidate early therapeutic target to modulate the potential for cyst formation.

Cell-cell communication analysis further revealed that SKiD cells act as active signaling hubs rather than mere recipients of disease cues. Both CNT and PT SKiD cells showed enhanced outgoing signaling, with overlap in NRG signaling. While previous reports have noted NRG pathway activation in both PKD (*62*) and acute kidney injury (*63*), the shift we observe from NRG4 signaling in control kidneys to tubular NRG1 signaling in PKD kidneys suggests a rewiring of epithelial communication networks during precystic disease. Importantly Nrg1 is a cleavable ligand, and its progressive accumulation specific to PKD kidneys identifies it as a promising non-invasive biomarker for early PKD progression, potentially enabling diagnosis or therapeutic intervention before cysts are sonographically apparent.

Together, these findings establish that PKD begins as a signaling disorder long before structural abnormalities emerge. SKiD cell cytoskeletal remodeling and CREB signaling provide a mechanistic framework for understanding how Pkd1 dysfunction primes kidneys for cystogenesis. This framework opens up new opportunities for early intervention including targeting CREB5-mediated CREB signaling and leveraging early biomarkers of disease such as NRG1. By defining increasingly earlier molecular events in PKD, this work lays the foundation for shifting our thinking of PKD from a cyst-centric disease to a molecular genetic disorder where we may exploit precystic signaling for disease prevention during a window when the kidney remains structurally intact and potentially most responsive to therapies.

## Materials and Methods

### PKD Mouse Model

All animal experimental procedures were conducted following the Seattle Children’s Research Institute’s Animal Care and Use Committee (IACUC) guidelines and procedures. The *Pkd1^RC/RC^*mouse model has been previously described and is a knock-in functionally hypomorphic model of PKD that results in gradual cystogenesis (*12*). Mice were maintained on a C57BL/6NJ background. Timed matings were conducted in the afternoon, and plugs were checked the following morning. Wild-type littermate mice were used as controls. Mouse kidneys were extracted at P5, P10, and P20. The one kidney per mouse was decapsulated and snap-frozen in liquid nitrogen for single nucleus transcriptomic analysis while the contralateral kidneys were fixed for histological and immunofluorescence studies.

### sci-RNA-seq3 Library Preparation and Sequencing

Library preparation was carried out by the Brotman Baty Institute Advanced Technology Lab (BAT-Lab) at the University of Washington. Each kidney was manually dissected with a scalpel in ice-cold lysis buffer (10 mM Tris-HCl, pH 7.4, 10 mM NaCl, 3 mM MgCl2, 0.1% IGEPAL CA-630, 1% SUPERase In, and 1% BSA). Kidney tissue was further manually homogenized with the rubber tip of a syringe plunger in 4 mL of lysis buffer prior to straining through a 40 μm cell strainer (Falcon). The strained nuclei were then transferred to a new 15 mL tube (Falcon) and pelleted by centrifugation at 500 G for 5 min and washed once with 1 ml cell lysis buffer. The nuclei were fixed in 4 ml of 4% paraformaldehyde (Electron Microscopy Sciences) for 15 min on ice. Following fixation, nuclei were washed twice in 1 mL wash buffer (lysis buffer without IGEPAL) and re-suspended in 500 μL wash buffer. Nuclei were permeabilized with 0.2% TritonX-100 (in wash buffer) for 3 min on ice, and briefly sonicated (Diagenode, 12 sec on low power mode) to reduce nuclei clumping. Nuclei were then washed once with wash buffer and strained through a 1 mL cell strainer (Flowmi). Nuclei were then centrifuged at 500 G for 5 min and resuspended in wash buffer.

One library was generated per kidney using the optimized sci-RNA-seq3 protocol (*64*). Nuclei were counted using the ImageExpress Pico System (Molecular Devices) and 15,000-20,000 nuclei were loaded per reverse transcription well. Tagmentation used N7 adaptor-loaded Tn5 transposase (Diagenode), with 6-9 μL of Tn5 per 550 μL of 2X TD buffer (20 mM Tris-HCl pH 7.6, 10 mM MgCl_2_, 20% dimethylformamide), distributed at 5uL per well. Libraries were cleaned using a two-step bead purification: first with 0.8X AMPure XP beads to remove lower molecular weight fragments, followed by a 0.55X bead-based supernatant step to remove larger fragments. Final purification was performed using GeneRead Size Selection columns (Qiagen). Libraries were sequenced using Illumina NovaSeq S4 flow cells at the University of Washington Northwest Genomics Center (settings: Read 1 - 34 cycles, Read 2 - 100 cycles, Index 1 – 10 cycles, Index 2 - 10 cycles).

### Cell clustering and data visualization

snRNA-seq preprocessing, read alignment, and gene count matrix generation was performed using the BAT-Lab pipeline (https://github.com/bbi-lab/bbi-dmux; https://github.com/bbi-lab/bbi-sci). This converts base calls to fastq files with bcl2fastq/ v.2.20 (RRID:SCR_015058), removes polyA tails using Trim Galore/v.0.6.7 (RRID:SCR_011847) (*65*), aligns trimmed reads to a reference genome with STAR/v.2.7.6 (RRID:SCR_004463) (*66*), extracts mapped reads, removes duplicates, and generates UMI counts for exonic and intronic regions of each gene, tabulated according to the unique three-level barcode design in sci-RNA-seq3.

The mouse reference genome (GRCm38/mm10) and Ensembl (version 101) (RRID:SCR_002344) annotation were used as reference. The 3’ untranslated region annotations of genes and transcripts were extended by 500 bp to avoid misclassifying genic reads as intergenic. After generating the count matrix, we removed nuclei of low quality or ambient RNA with the following standards: 1. UMI counts < 100, 2. number of gene detected ≤ 100, 3. percentage of mitochondria ≥ 10%, and 4. percentage of reads mapping to intronic region ≥ 20%. For each sample, we then imported the filtered gene-by-nucleus count matrices into the AnnData/v.0.8.0 (RRID:SCR_018209) (*67*) framework and then used Scrublet/v.0.2.3 (RRID:SCR_018098) (*68*) to calculate doublet scores. We marked nuclei as Scrublet-inferred doublets if they had Scrublet doublet scores > 0.20.

Downstream analyses were performed with Monocle3 (RRID:SCR_018685) (*19*, *69*). We visualized the filtered dataset by UMAP. We annotated each cell to its putative cell type manually based on marker gene expression. After finalizing the annotation of the cell type in each cluster, we combined nuclei of the same cell types to generate subclusters. We performed normalization and dimensionality reduction as above for each subcluster. The subcluster enrichment threshold of 66% of cells derived from a single-condition was based on the relative proportion that would represent a statistically significant increase that is not readily explained by variable cell sampling as predicted by multiple models (*70*).

### Differential Gene Expression Analysis

Differential gene expression analyses were carried out across all annotated cell types. A quasi-Poisson distribution model was used to analyze gene expression data comparing nuclei originating from WT to PKD kidneys. We performed false discovery rate (FDR) correction on the resulting p-values and considered genes significant at an adjusted p-value < 0.05. We generated graphs of differential gene expression to highlight genes meeting this significance threshold along with a fold expression change of < log_2_(-1.5) or > log_2_(1.5).

### Pathway Analysis

Pathway analyses were carried out to compare subclusters within annotated cell types with PKD-enriched subclusters. PKD-enriched subclusters were defined as subclusters in which more than 66% of nuclei originated from PKD kidneys. For each cell type with a PKD-enriched subcluster we imported the entire cell type dataset into VISION/v.3.0.2 (*20*). Enrichment analysis was conducted using BioCarta Pathways (RRID:SCR_006917) accessed through the Molecular Signature Database (RRID:SCR_016863) (*71*).

### Pseudotime Analysis

We performed normalization and dimensionality reduction of a dataset consisting of nuclei annotated as DCT or PT across all timepoints. The pseudotime analysis was performed with Monocle3 using default parameters.

### Module Analysis

To identify coregulated gene modules, using Monocle3 we used UMAP dimensional reduction to cluster genes based on similar expression levels across a dataset containing nuclei annotated as DCT or PT across all timepoints. We subsequently grouped them into modules using Louvain community analysis. We then grouped modules both by timepoint and subclustered identity to identify modules enriched at times and cell subtypes potentially relevant to PKD.

### Transcription Factor Analysis

We queried publicly available datasets in the NCBI Gene Expression Omnibus to identify gene sets from experiments in which a transcription factor was overexpressed or disrupted (e.g. by knock-out, knock-down, or dominant negative approaches) as previously reported (*72*). We assessed overlapping gene sets with these experiments to the gene sets represented in our identified gene modules.

### Immunofluorescence Labeling and Microscopy

Five-micron thick coronal sections of mouse kidney were prepared from paraffin embedded kidneys of either WT or PKD mice at P5, P10, or P20. Slides were heated to 60 °C for 2 hours prior to antigen retrieval in 1 mM EDTA pH 8.0 at 95 °C for 15 min. Sections underwent deparaffinization through a series of xylene and ethanol washes for a duration of 3 min each (xylene #1, xylene #2, xylene #3, 100% ethanol #1, 100% ethanol #2, 90% ethanol, 75% ethanol, 50% ethanol, and deionized water).

Immunofluorescence labeling of proteins was completed using antibodies at the concentrations listed in **Supplementary Table 1**. Briefly, sections were blocked overnight in 10% FBS in PBS + 0.1% Tween-20. Sections were then incubated in primary antibody in 10% FBS in PBS + 0.1% Tween-20 for 2 h and rinsed twice with PBS and washed twice in PBS for 15 min. Sections were then incubated in secondary antibody in 10% FBS in PBS + 0.1% Tween-20 for 1 h. Sections were then incubated in 10 mg/mL Hoechst 3342 (Sigma-Aldrich) for 15 min and rinsed twice with PBS and washed twice in PBS for 15 min.

Prior to imaging, samples were mounted in Prolong Glass medium (Thermo Fisher Scientific) and covered with #1.5H cover glass. Labeled kidney sections were imaged using a Zeiss LSM 900 microscope with Airyscan 2. Lasers with wavelengths of 405, 488, 561, and 640 nm were used for excitation. Images were acquired using a 20x Plan Apochromat air objective. The images were captured sequentially, with a pixel dwell time of 2 µs and a 1 AU pinhole without averaging. Zeiss Zen 3.12 acquisition software was used during imaging. Fluorescent micrographs were processed and analyzed with Fiji (*73*). PT cells were identified based on positive LTL labeling. CNT cells were identified based on the combination of antibody based detection of Slc8a1 and absence of antibody based detection of Slc12a3.

### Cell-Cell Communication Analysis

Cell-cell communication ligand-receptor analysis was performed with CellChat v2.1.2 (RRID:SCR_021946) (*54*). Two CellChat objects were generated using 1: all WT PT and DCT clusters across P5, P10, and P20 and 2: all WT PT and DCT clusters across P5, P10, and P20. Following standard preprocessing and computation of communication probability, cell-cell communication was compared between datasets using compareInteractions functions. Data visualization was completed with netVisual_diffInteraction, netVisual_bubble, and netVisual_chord_gene functions.

### Urine Collection

Mice were removed from their enclosures and the lower abdomen above the level of the bladder was slowly rubbed with a gloved finger with minimal pressure applied toward the urethral meatus to induce urination.

### Immunoblotting

Urine samples from individual mice were combined with 10% 2-mercaptoethanol containing Laemmli buffer and boiled at 95 °C for 5 min. The samples were loaded into the wells of 4–20% Mini-PROTEAN® TGX Stain-Free Protein Gels (Bio-Rad). Electrophoresis and transfer were completed through standard methods. Immunoblots were blocked overnight in blocking buffer (LICORBio). Blots were then incubated in primary antibody in blocking buffer for 2 h and rinsed twice with TBS and washed twice in TBS for 15 min. Blots were then incubated in secondary antibody in blocking buffer for 1 h and rinsed twice with TBS and washed twice in TBS for 15 min. Blots were imaged using an Odyssey F imaging system (LICORBio).

## Data Availability

The data generated by this study can be downloaded in raw and processed forms from the NCBI Gene Expression Omnibus.

## Code Availability

The code used to analyze data as part of this study are publicly accessible (github.com/MARQUEZ-PKD/RC).

## Author Contributions

Conceptualization: J.M., E.D.N., and D.R.B. Methodology: J.M., O.T., G.H., and A.M.M. Investigation: J.M., O.T., S.K. G., S.R.H., G.H., A.M.M., and E.D.G. Formal Analysis: J.M., E.D.N. and D.R.B., Writing–original draft: J.M., E.D.N. and D.R.B., Writing–review & editing: all authors.

## Data and Materials Availability

All data needed to evaluate the conclusions in the paper are present in the paper and/or the Supplementary Materials. The sequencing data generated in this study have been deposited in the GEO database.

We thank the Brotman Baty Institute advanced technology lab for technical assistance in completing single nucleus RNA-sequencing. We thank the Seattle Children’s Research Institute Microscopy and Histopathology Collaborative Laboratory for access to microscopy resources. We thank the staff of the Office of Animal Care for assistance in housing and caring for mice used in this study. We thank Peter C. Harris of the Mayo Clinic Robert M. and Billie Kelley Pirnie Translational Polycystic Kidney Disease Center for initial provision of the RC mouse model.

