## Supplemental Figures for "Single-nucleus Transcriptomics Reveals Precystic Dysfunction in Polycystic Kidney Disease"

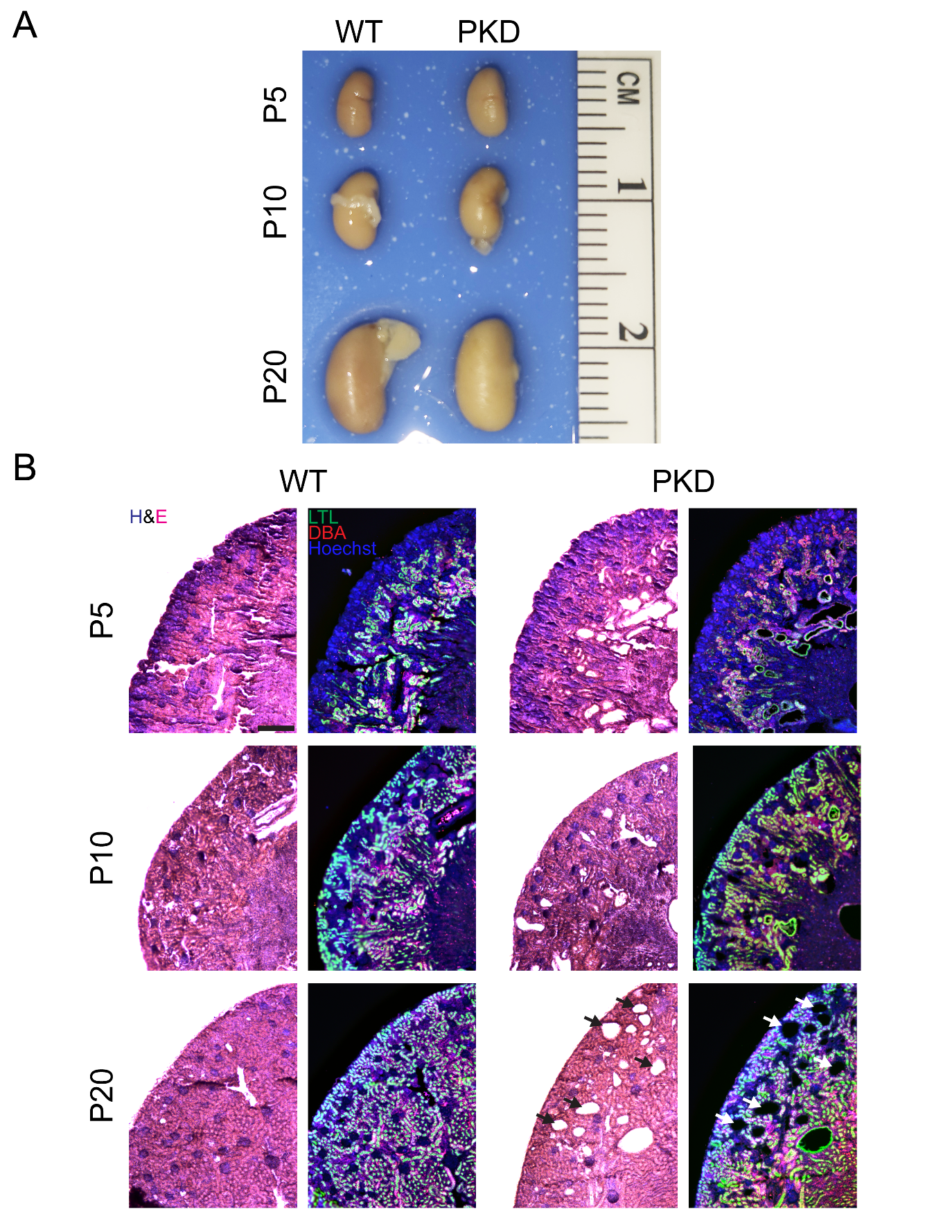

**Supplementary Figure 1: Progression of cyst formation in RC mouse model of PKD** (A) Image of extracted WT and PKD kidneys. (B) Hematoxylin and eosin staining along with fluorescence imaging demonstrate little cystic tissue at P5 or P10 in either WT or RC kidneys though cysts identifiable in P20 RC kidneys (arrows).

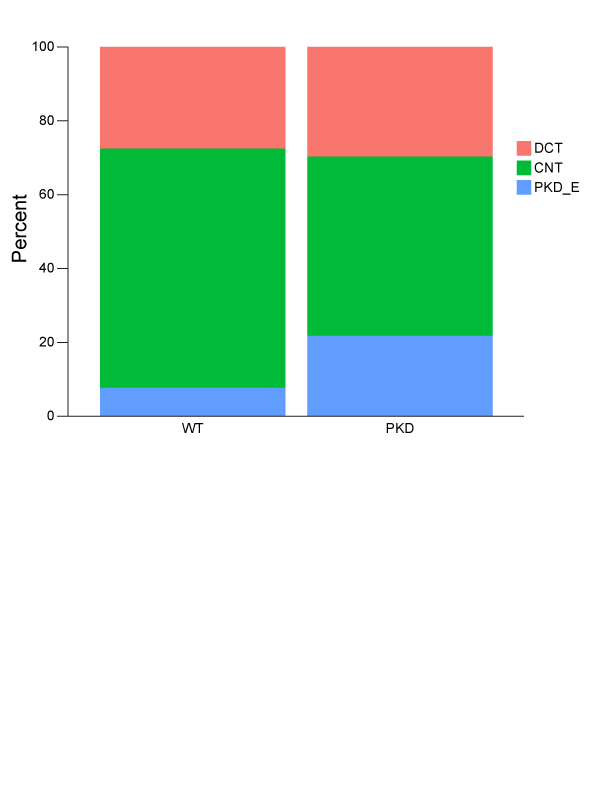

**Supplementary Figure 2: Proportion of nuclei within DCT subclusters**

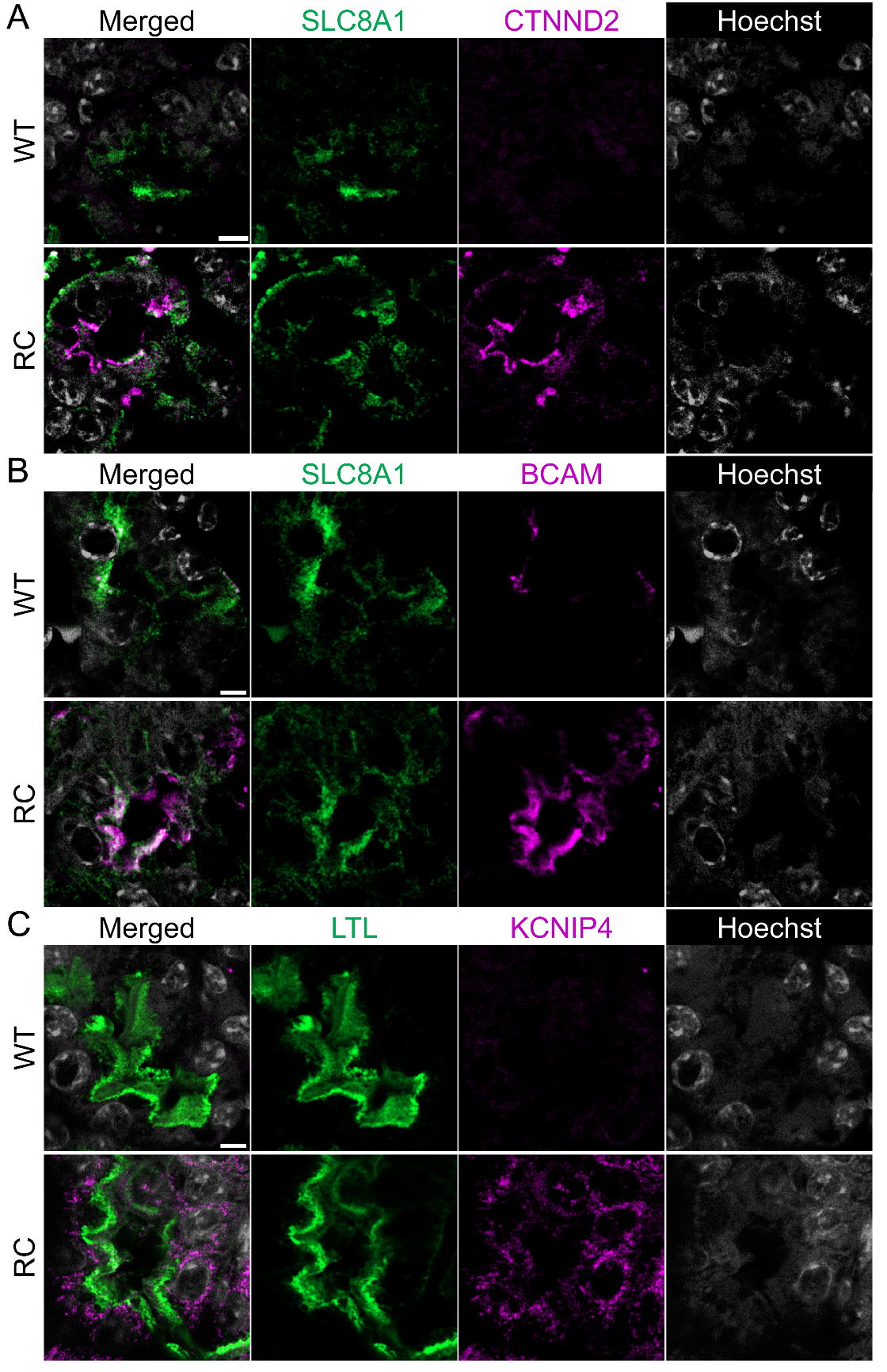

**Supplementary Figure 3: Detailed Immunofluorescence Assessment of Proteins Encoded by Genes Upregulated in PKD Enriched Subclusters** (**A**) Immunofluorescent labeling of CTNND2 in WT and PKD CNT cells at P20. (**B**) Immunofluorescent labeling of BCAM in WT and PKD CNT cells at P10. (**C**) Immunofluorescent labeling of KCNIP4 in WT and PKD PT cells at P10.

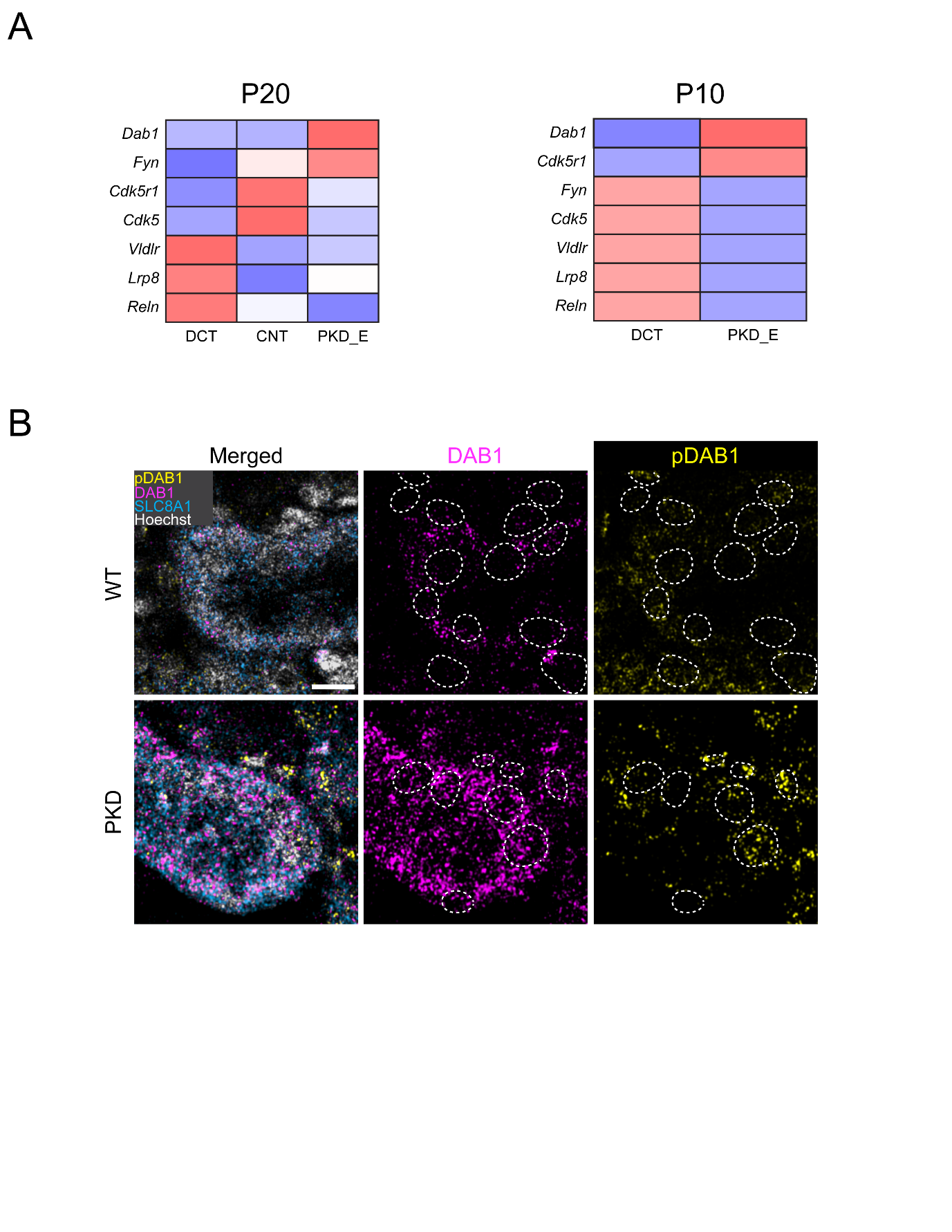

**Supplementary Figure 4: Non-canonical REELIN signaling Is Increased in Precystic PKD Kidneys** (A) Signature heatmap of REELIN signaling components across P10 DCT subclusters. (B) Comparison of DAB1 and pDAB1 abundance in WT and PKD kidneys at P10. Scale bar indicates 5 μm.

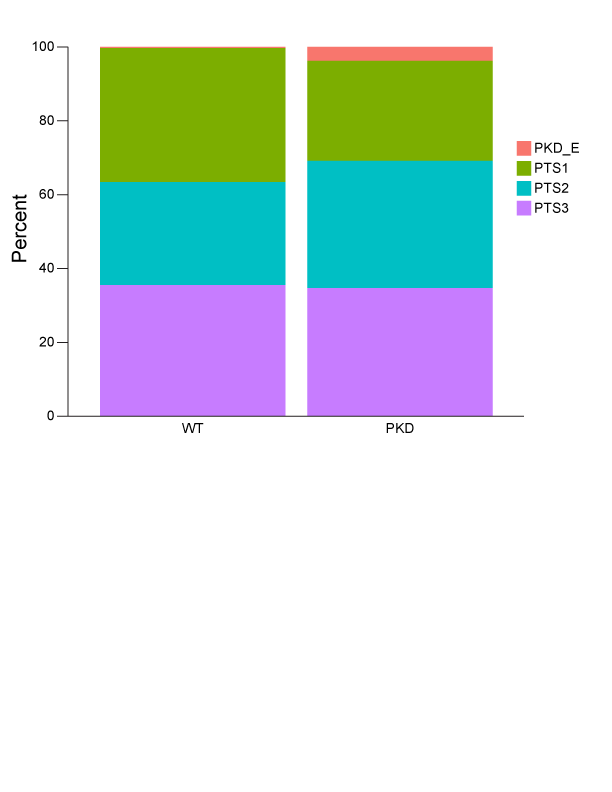

**Supplementary Figure 5: Proportion of nuclei within PT subclusters**

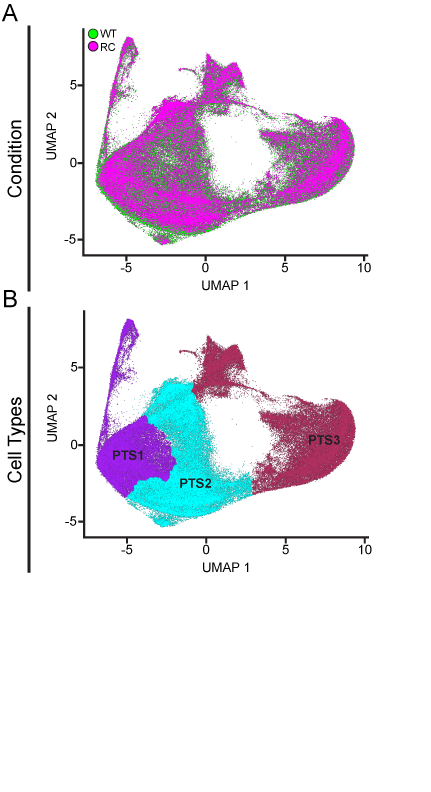

**Supplementary Figure 6: A PKD Enriched Cell State is Not Apparent in P20 PT Cells** (**A**) UMAP plot of subclustered snRNA-seq dataset of P20 PT cluster colored by tissue of origin genotype. (**B**) UMAP plot of subclustered snRNA-seq dataset of P20 PT cluster colored by subcluster cell type annotation.

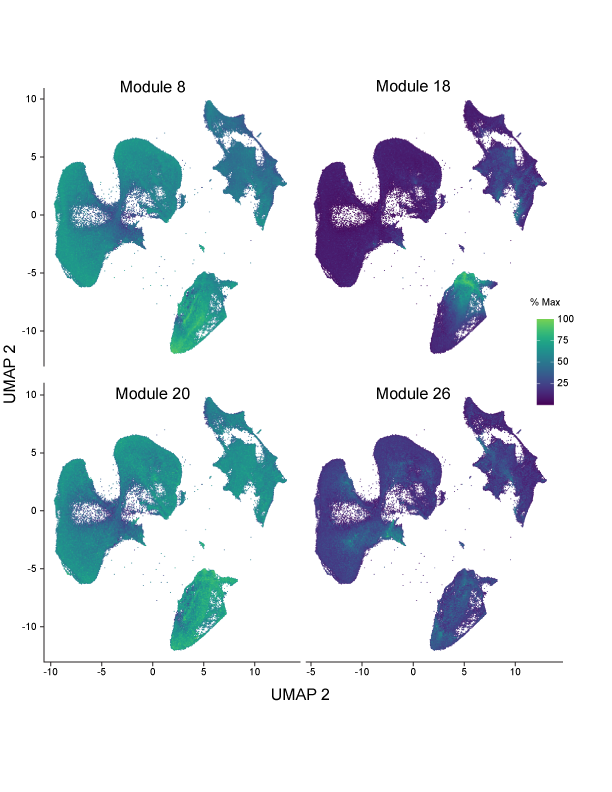

**Supplementary Figure 7: Modules of Coregulated Genes Show Increased Expression in PKD Relevant Nuclei**

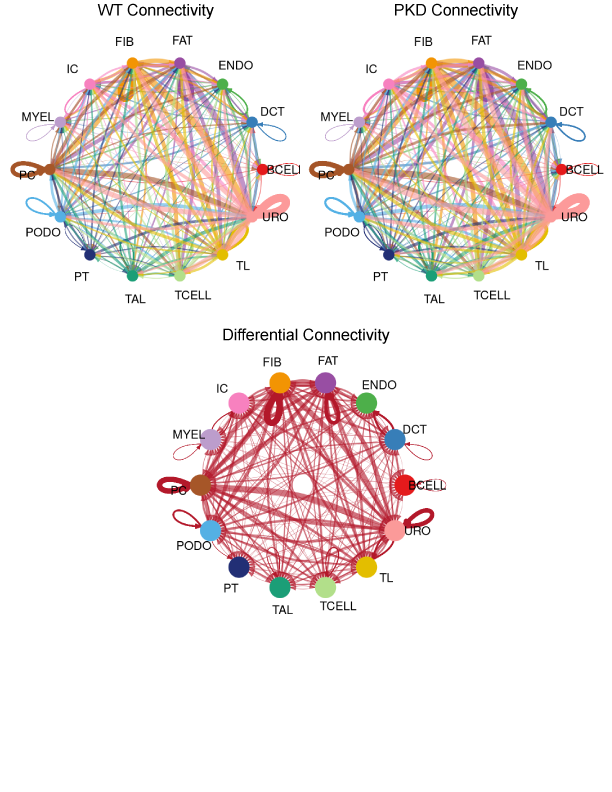

**Supplementary Figure 8: String diagram of differential interaction strength between all annotated cell clusters between WT and PKD nuclei in P10 kidneys.** Arrow width is proportional to increased communication in PKD tissue.

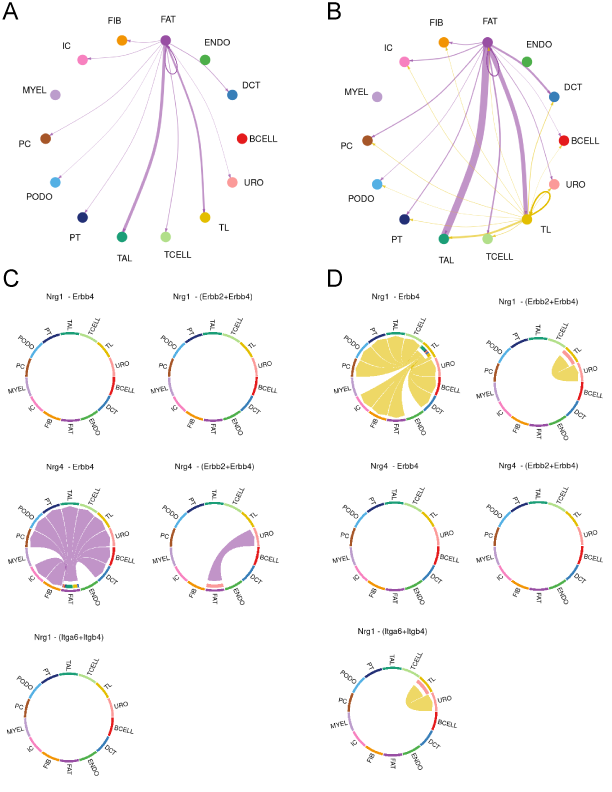

**Supplementary Figure 9: NRG Pathway Connectivity Demonstrates Distinct Patterns in WT and PKD contexts** (**A**) String diagram of NRG pathway cell-cell communication in WT kidneys. (**B**) String diagram of NRG pathway cell-cell communication in PKD kidneys. (**C**) Chord plot of ligand-receptor pairs mediating NRG pathway cell-cell communication from fat cells in PKD kidneys. (**D**) Chord plot of ligand-receptor pairs mediating NRG pathway cell-cell communication from thin limb cells in PKD kidneys.

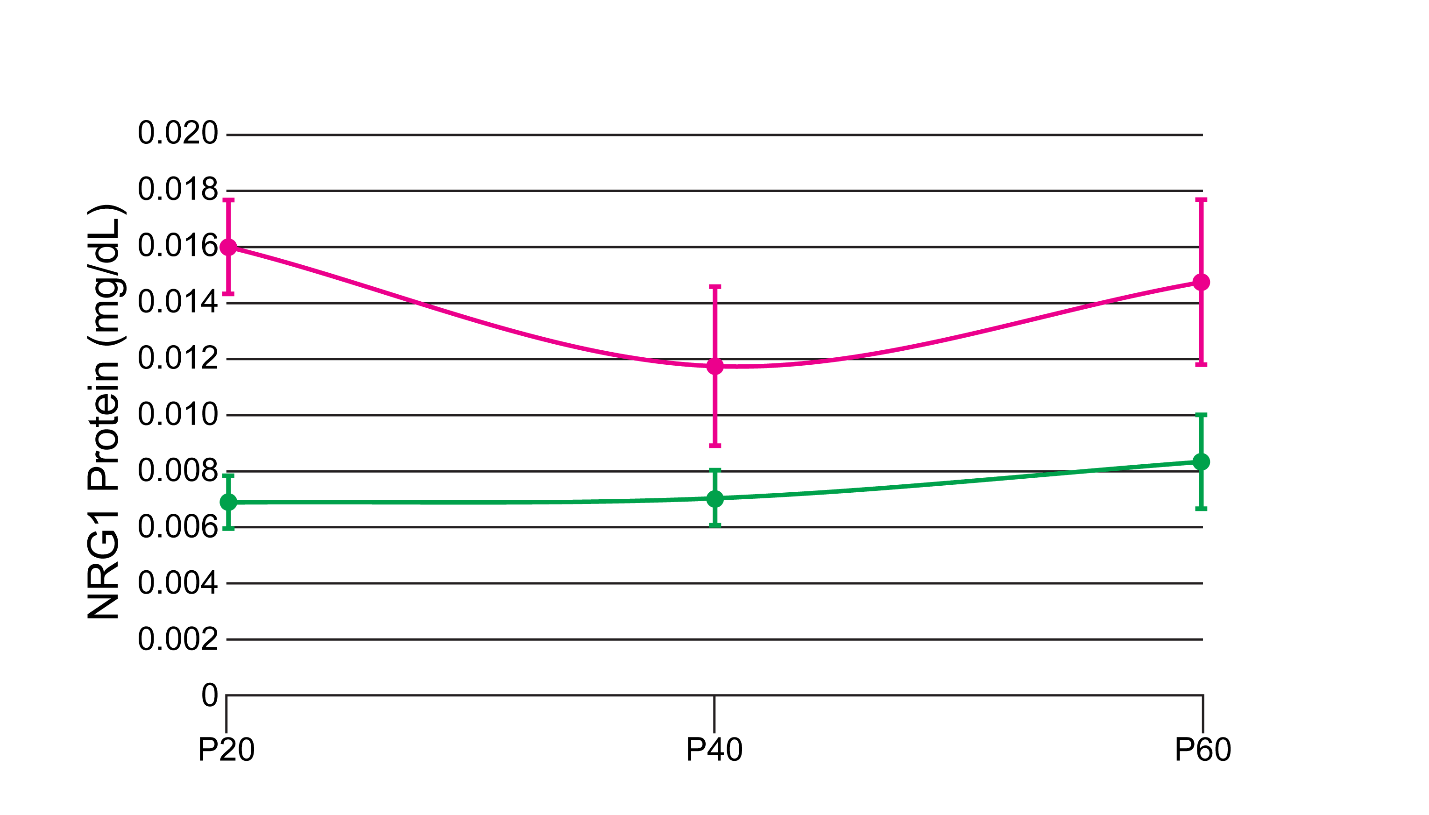

**Supplementary Figure 10: NRG1 is Detectable in Urine and is Increased in PKD Mice** Quantitation of urine Nrg1 levels in WT and PKD mouse urine. Dots represent means with vertical bars representing standard deviation.
